# Evolutionary History Impacts Phyllosphere Community Assembly on Forage Grasses

**DOI:** 10.1101/2021.06.15.448595

**Authors:** Emily K. Bechtold, Klaus Nüsslein

**Author notes:** Funding: The efforts of Emily Bechtold in this work were funded by the Lotta M. Crabtree Foundation. Additional research support was provided by the National Science Foundation – Dimensions of Biodiversity (DEB 1442183).

## Abstract

Benefits leaf bacterial communities provide to plant hosts are reduced by external stress. Understanding how plant hosts impact phyllosphere community assembly, how microbes influence plant traits, and how this interaction changes under stress will advance our insight into the evolutionary relationship between plants and their microbial communities. We investigated phyllosphere community assembly change over time, between host species, and under drought stress on three native temperate grasses and three non-native tropical grasses. By growing them together, effects of host geography and differences in environmental variables were eliminated allowing us to test evolutionary history on community assembly. We found evidence of phylosymbiosis which increased significantly under drought stress, indicating phyllosphere communities and their response to stress relate to grass species phylogeny. We also show native temperate grasses displayed stronger cophylogenetic relationships between grass hosts and their microbial communities and had increased selection by host species over time compared to non-native tropical hosts. Interestingly, the functional marker gene *nifH*, though differentially present on all host species was not susceptible to drought. The evidence of shared evolutionary history, presence of functionally important bacteria, and responses to drought suggest that microbial communities are important plant traits that coevolve alongside their plant hosts.

## INTRODUCTION

As one of the largest terrestrial habitats, grasslands make up nearly 70% of global agricultural land and contribute important ecosystem services including impacting water quality, erosion prevention, and climate regulation through carbon sequestration and greenhouse gas mitigation [1]. The important agricultural and ecological functions grasslands provide are threatened due to projected decreases in water availability as drought frequency and severity continue to increase [2–4]. This will have drastic effects on grassland productivity, ultimately reducing global food security and increasing climate change [5].

Grass leave surfaces harbor diverse microbial communities, termed the phyllosphere, which provide important functions to their host including disease prevention, stress tolerance, ecosystem productivity, and nutrient cycling through processes such as nitrogen fixation [6–9]. In return, plants provide nutrients to the bacteria creating a symbiotic relationship, but what drives these relationships is not completely understood. Previous studies show that while plant host identity plays an important role in microbial community assembly, phyllosphere communities are broadly dominated by similar taxa including *Proteobacteria, Bacteroidetes*, and *Actinobacteria* [10–13].

Common theories to explain phyllosphere community assembly include the existence of a functional core community, in which phyllosphere community members provide consistent functional traits across host species [11, 14], and the hologenome theory of evolution, which postulates evolution occurs between hosts and microbes together [15]. Many core functions support epiphytic bacterial growth under harsh conditions indicating microbial adaptation to the phyllosphere. These include pigmentation and DNA repair systems to protect from UV radiation production of extracellular polysaccharides to promote biofilm formation which protects against osmotic stress, and motility-related proteins for movement towards nutrients [8, 11, 16, 17]. Additional functional traits are important for plant health and ecosystem functioning, by promoting global carbon and nitrogen cycles, photosynthetic strategies, resource acquisition, and plant defense [11, 14, 18]. Nitrogen fixation by bacteria called diazotrophs, frequently associated with the rhizosphere, occurs in the phyllosphere contributing to total nitrogen input in an ecosystem [9, 11, 19]. The observed differences in relative abundance of functional genes and taxonomic identity despite low variability between host species [13, 14], suggest the functional core exists within the hologenome theory. For example, phyllosphere bacteria can have rhodopsins which provide energy and protection from UV damage. These pigments absorb different wavelengths of light than their plant host allowing for optimal utilization of resources thus indicating shared evolutionary history [20–22].

How functional profiles and plant-microbe relationships change under stress conditions is still unknown. Therefore, we do not understand if response to stress is a stochastic process dependent largely on atmospheric conditions, a response to changes in plant physiology, or a response characterizing joint plant-microbe interactions. One method to explore host-microbe relationships is phylosymbiosis, which determines if significant associations between microbial communities and the phylogeny of their host species exist [23–25]. Phylosymbiosis can be determined using a Mantel test to compare a host phylogenetic distance matrix to a microbial community distance matrix. When phylosymbiosis occurs, phylogenetically related host species have more similar microbial communities than less phylogenetically related hosts. Phylosymbiosis can result from coevolution, which occurs when plant-microbe systems act as reciprocal selective forces on each other [25, 26]. However, it can also result from differences in host geography, host traits, or codiversification, which occurs when hosts and microbes exhibit parallel divergence during continued associations [27]. A second method used to understand how host phylogeny relates to microbial communities is cophylogeny, which tests the concordance of the host phylogeny with the phylogeny of the associated microbial community [28, 29]. Cophylogenetic occurrences indicate shared evolutionary history between hosts and microbial groups [29, 30]. While cophylogeny can result from processes such as biogeographical distance, presence of cophylogeny is consistent with host-microbe coevolution [29, 31, 32]. Previous work suggests that cophylogenetic associations are more likely to exhibit microbe-to-host interactions [14, 33, 34]. Therefore, identifying these associations can help identify evolutionarily important and ecologically active plant-microbe relationships.

To understand plant-microbe interactions we need to understand rules of assembly and functional processes. Our objective was to investigate if phyllosphere communities are an adapted plant trait. To address this objective, we explored the questions: (i) How does host phylogeny influence microbial community assembly? (ii) How does host identity or phylogeny influence microbial community response to drought stress? (iii) How is diazotroph abundance related to microbial community structure and response to stress? To answer these questions, we investigated how microbial community assembly changed over time, between host species, and under drought stress. We chose three species of grasses commonly used in temperate forage systems and three species commonly used in tropical forage systems. By growing all species in the same common garden experiment, we eliminated effects of host geography and differences in environmental variables on community assembly. Additionally, by growing native temperate and foreign tropical species, we tested the influence of evolutionary history on community assembly. Comparing the evolutionary history of phyllosphere communities to that of their hosts and determining how communities change under drought stress, allowed us to understand if phyllosphere microbes are a plant trait and begin to understand how to leverage microbes to promote plant growth and stress tolerance.

## MATERIALS AND METHODS

### Study system

Seeds for three non-native tropical grasses, *Brachiaria brizantha* (CIAT 26564), *Brachiaria decumbens* (CIAT 6370), and *Brachiaria* hybrid (CIAT 1794), were acquired from CIAT (Cali, Columbia). Native temperate grass species seeds, *Festuca arundinacea* (endophyte free Tall Fescue), *Dactylis glomerata* (Orchardgrass), and *Lolium perenne* (Ryegrass), were acquired from Albert Lea Seed Company (Albert Lea, MN, USA). Seeds were germinated in Pro-mix commercial potting medium (Quakertown, PA, USA) in 2018 and grown in the College of Natural Sciences Research and Education Greenhouse at the University of Massachusetts-Amherst. In June 2019, individual plants were transplanted into 15×30cm pots filled with soil collected from natural grass fields in Amherst, MA (Supplementary Methods, Supplementary Table 1). Pots were moved outside, organized in a randomized block design, and allowed to re-establish. Ten plant replicates of each temperate species were divided between ‘control’ and ‘drought’ treatments. Drought treatment plants were placed under a 10 ft high rain shelter made of greenhouse plastic allowing maximal airflow and high UV light penetration (Supplementary Figure 1). Drought conditions were imposed over 38 days (21 AUG - 27 SEPT 2019). Plants in the control group were given supplemental water to maintain soil moisture above 80% field capacity. Plants in the drought group were given supplemental water when necessary to maintain an even dry-down rate, determined from soil-moisture readings measured twice weekly using a MiniTrase TDR with Buriable probe (Soilmoisture Equipment Corp., Goleta, CA, USA).

### Plant Health Measurements

Plant health measurements were taken on days 1, 19, 26, 33, and 38 to understand the effect of drought on the plant host. Plant measurements taken were leaf relative water content (RWC), chlorophyll concentration, and leaf cellular membrane stability determined by measuring electrolyte leakage [35–37]. At the end of the drought period, above ground biomass was measured by dividing plant material into five categories: stems, flowers, dead, mature, and young leaves. After determining fresh mass, samples were dried in an incubator at 70°C for 5 days and dry mass was measured.

### Bacteria Community Sampling

At each timestep, bacterial community DNA was extracted using the Nucleospin Plant II Extraction Kit (Machery-Nagel, Düren, Germany) following a modified protocol. Five whole ryegrass leaves or three whole leaves of each other species were aseptically removed from the plant host and placed into a 15 ml conical tube with 1.5 ml of NucleoSpin Type-B beads and 4X volume of Buffer PL1. Tubes were vortexed horizontally for 5 min at room temperature. The lysate was incubated for 60 min at 65°C, placed in a NucleoSpin Filter tube, and centrifuged for 2 min at 11,000x*g*. The filtrate was added to 4X Buffer PC and extraction continued following the manual. Aydogan et al. found that vortexing whole leaf samples in tubes with lysis buffer and beads extracted important community members from biofilms with minimal plant DNA co-extraction [12]. Extracted DNA samples underwent a two-step PCR amplification to attach Illumina adaptor sequences and barcodes (Supplementary Methods). The first PCR step used chloroplast excluding primers 799F and 1115R targeting the V5-V6 region of the 16S rRNA gene [10] with linker sequences to attach Access Array Barcodes (Fluidigm, San Francisco, CA, USA) [38]. Amplicons were pooled and sequenced on Illumina MiSeq Platform, with 251 bp paired-end sequencing chemistry at the Genomics Resource Laboratory (University of Massachusetts-Amherst). The abundance of nitrogen-fixing bacteria was determined using qPCR quantifying the *nifH* gene using the PolF and PolR primers [39].

### Sequence Analysis

Using the QIIME2 [40] pipeline, paired-end reads were demultiplexed, merged, trimmed to 315 base pairs, and binned inferring amplicon sequence variants (ASVs). Taxonomic identities were assigned using the naïve Bayes sklearn classifier trained with the 799F/1115R region of the Greengenes 13_8 database.

The data contained 9,207 ASVs from 280 samples containing a total of 15,218,029 reads. Samples were rarefied to 4,000 reads, resulting in a loss of 16 samples. Alpha diversity was calculated using Shannon Diversity Index and beta diversity using Weighted UniFrac and Bray-Curtis distance metrics.

### Machine learning

We used the mikropml R package to conduct machine learning (ML) analyses [41–43]. For each model, we used random forest classification with 75% of the test data used to train the model and the remaining 25% to test the model. ML was used on data collected the last day of drought to predict if: (1) communities are from control or drought treated plant hosts regardless of host species, and (2) bacterial communities came from tropical or temperate grass hosts regardless of treatment. Model performance was evaluated using the area under the operating characteristic curve (AUC) value. Models yielding AUC values above 0.6 were determined to have good predictive power. Additionally, the mikropml pipeline enables determination of bacterial features important for prediction and how much they contribute to AUC values.

### Phylosymbiosis and Cophylogeny

Phylosymbiosis was determined using a Mantel test with matrices of grass species’ phylogenetic distances and microbial community beta diversities calculated using Bray-Curtis and weighted UniFrac distances. Grass host phylogenetic distances were calculated using MEGAX [44]. Sequences of the chloroplast gene for each species were retrieved from NCBI [45] and aligned using MUSCLE [46]. A phylogenetic tree was constructed using the maximum likelihood method. A Mantel test was performed with the Spearman’s Rank correlation with 9999 permutations using the Vegan package in R [47].

We tested for cophylogeny to understand if coevolution between microbial communities and their plant host exists. Two separate global fit methods were employed: ParaFit as carried out in the ape package [48] and PACo using the R package paco [30]. Microbial data used to test for cophylogeny were filtered to only include data collected on the last sampling day with at least 100 reads across all samples resulting in 359 ASVs. Both methods were performed with host phylogeny, microbial 16S rRNA phylogeny, and a presence/absence matrix for each host and ASV. Additionally, both methods used the Caillez correction method to account for negative eigenvalues. PACo analysis was performed with 1000 permutations using the most conservative quasiswap method, which is used when it is uncertain if the host is tracking symbiont evolution or symbionts are tracking host evolution. ParaFit was performed using 999 permutations. Significant associations were plotted using the cophyloplot function in the ape package.

### Statistical methods

Separate generalized linear mixed models (GLMMs) were created to assess changes in alpha diversity and *nifH* abundance using gamma distributions with a log link using the lme4 R package [49]. Drought treatment, host species, and time were fixed effects and sample ID a random effect to account for sampling over time. Effects of each variable was determined using Tukey tests for comparison using lsmeans [50]. Effects of host species, drought treatment, time, and their interactions on microbial community structure were determined using permutational analysis of variance (PERMANOVA) and analysis of multivariate homogeneity of group dispersions (PERMDISP2) with weighted UniFrac distances. Results were visualized using non-metric multidimensional scaling (NMDS). PERMANOVA, PERMDISP2, and NMDS were conducted using the vegan package and visualized using ggplot2 [51].

## RESULTS

Phyllosphere communities varied between host species, over time, and as a result of drought. Across all sample days and host species, *Alphaproteobacteria* was the dominant class in both control (34.2%) and drought samples (34.6%), but community dynamics over time and as a result of drought were different between host species (Figure 1A). At the start of the experiment, *Alphaproteobacteria* and *Gammaproteobacteria* were the dominant groups, but by the end of the experiment *Cytophagia* was the dominant class under control conditions. While *Cytophagia* increased in relative abundance under drought conditions, *Alphaproteobacteria* remained the dominant group on drought stressed hosts.

**Figure 1.**
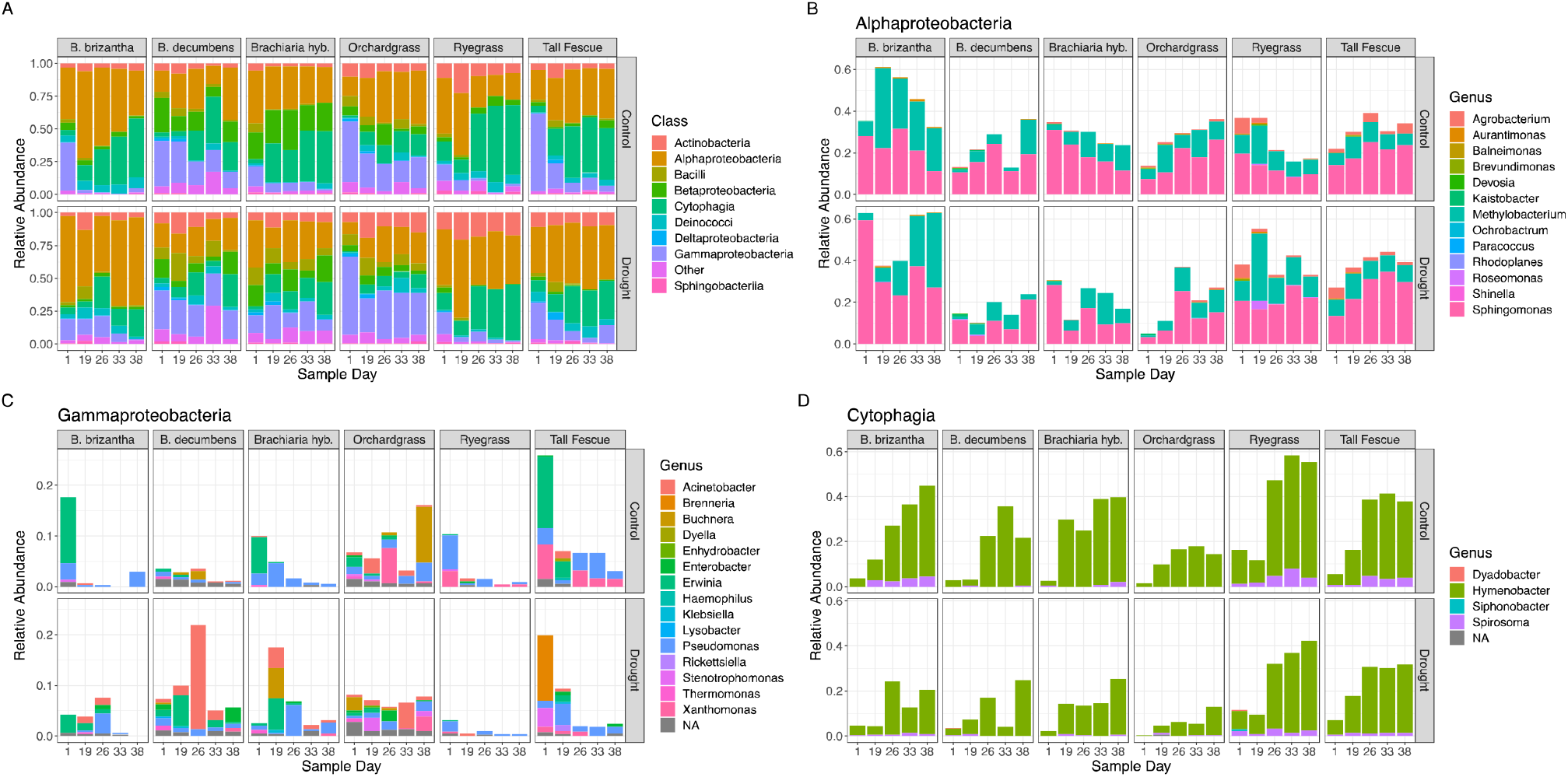
Average relative abundance of bacteria from 57 plants (27 control and 30 drought) sampled at 5 separate time points over 38 days. (A) The most dominant bacterial classes changed over time, between host species, and as a result of drought. To understand the composition of these classes, the average relative abundance of the genera from the three most abundant classes were plotted. Genera included were present in greater than 0.25% average relative abundance. At the end of 38 days when drought effect was strongest, we observed significant differences as a result of drought in *Actinobacteria* (P<0.001), *Bacilli* (P=0.006), and *Cytophagia* (P=0.001) (calculated using TukeyHSD). Additionally, strong differences were observed between host species with significant differences observed in *Actinobacteria* (P<0.001), *Alphaproteobacteria* (P<0.001), *Bacilli* (P<0.001), *Betaproteobacteria* (P<0.001), *Cytophagia* (P<0.001), *Deltaproteobacteria* (P=0.001), and *Gammaproteobacteria* (P=0.008) (B) The class Alphaproteobacteria was dominated by the genera *Sphingomonas* and *Methylobacterium*, (C) Gammaproteobacteria was not consistently dominated by any individual genera, and (D) the class Cytophagia was dominated by the genus *Hymenobacter*.

*Alphaproteobacteria* was dominated by *Sphingomonas* and *Methylobacterium* for each species, but trends in relative abundance over time and as a result of drought were different between the host species (Figure 1B). Genera from the class *Gammaproteobacteria* were more diverse and variable between treatments, host species, and over time, but *Pseudomonas* was consistently present across samples (Figure 1C). The increase in *Cytophagia* was accounted for almost exclusively by the genus *Hymenobacter* (Figure 1D).

### Host species, time, and drought drive changes in community structure

To evaluate the role plant species and drought had on microbial community diversity, we modeled how alpha diversity changed over time, as a result of drought, and based on host species (Supplementary Figure S2). Alpha diversity was not affected by drought treatment but was significantly different based on host species identity.

Phyllosphere community structures were impacted by time, host species, and drought. Additionally, the degree microbial communities changed as a result of drought related to known drought tolerances of their host species. The strongest driver of phyllosphere community structure was plant host species (R^2^= 0.19, p=0.00; PERMANOVA on weighted UniFrac distances) (Table 1). Sample day (R^2^=0.14, p=0.001) and watering condition (R^2^=0.02, p=0.001) were also significant drivers. All three two-way interactions were significant, with the strongest interaction between sampling day and host species (R^2^=0.10, p=0.001). However, the three-way interaction was not significant (R^2^=0.04, p=0.0572). PERMDISP2 was conducted to ensure significant PERMANOVA results were caused by shifts in community structure instead of differences in dispersion within treatments. PERMDISP2 analyses were not significant (p=0.07), indicating that significant results from the PERMANOVA analyses are important factors for community structure. Because of significant two-way interactions, we conducted individual analyses on host species and sampling day to understand how microbial communities from each host changed over time and were impacted by drought. Overall community response was first detected 33 days into the experimental period. Additionally, host species effect on community structure increased over time (Figure 2). Separate PERMANOVAs run on control samples from Day 1 (R^2^=0.38, p=0.01) and Day 38 (R^2^=0.57, p=0.001) show increased effect of host species on community assembly under non-stressed conditions. Additionally, influence of host species within the temperate (R^2^=0.34, p=0.039) and tropical (R^2^=0.31, p=0.094) groups was similar at the start of the experiment, but temperate species (R^2^=0.72, p=0.001) explained greater variability by the end of the experiment than the tropical species (R^2^=0.39, p=0.024) (Table 2).

**Table 1.** Phyllosphere community structure on native temperate and non-native tropical grasses change over time (day) and are impacted by host species and drought treatment. Impact of each variable on community structure was determined using a PERMANOVA on weighted UniFrac distance measures.

| Variable | Weighted UniFrac Distance |  |
| --- | --- | --- |
|  | **R<sup>2</sup>** | **P-value** |
| Treatment | 0.019 | 0.001*** |
| Day | 0.142 | 0.001*** |
| Species | 0.191 | 0.001*** |
| Treatment*Day | 0.017 | 0.008** |
| Treatment*Species | 0.029 | 0.001*** |
| Day*Species | 0.101 | 0.001*** |
| Treatment*Day*Species | 0.043 | 0.0572 |

**Table 2.** The effect of host species on phyllosphere community composition on non-stressed hosts increased over time. The impact of host species on community structure was measured for communities from the well-watered control host plants at (A) the beginning (day 1) and (B) the end (day 38) of the drought period. Influence of host species was determined for all hosts together and separately for the native temperate grasses and non-native tropical grasses using a PERMANOVA of weighted UniFrac distances.

| DAY 1 | All host plants |  | Temperate |  | Tropical |  |
| --- | --- | --- | --- | --- | --- | --- |
|  | **R<sup>2</sup>** | **P-value** | **R<sup>2</sup>** | **P-value** | **R<sup>2</sup>** | **P-value** |
| Species | 0.38 | 0.01 | 0.34 | 0.039 | 0.31 | 0.094 |
| Residuals | 0.62 |  | 0.65 |  | 0.69 |  |

B.
| DAY 38 | All host plants |  | Temperate |  | Tropical |  |
| --- | --- | --- | --- | --- | --- | --- |
|  | **R<sup>2</sup>** | **P-value** | **R<sup>2</sup>** | **P-value** | **R<sup>2</sup>** | **P-value** |
| Species | 0.57 | 0.001 | 0.72 | 0.001 | 0.39 | 0.024 |
| Residuals | 0.43 |  | 0.28 |  | 0.61 |  |

**Figure 2.**
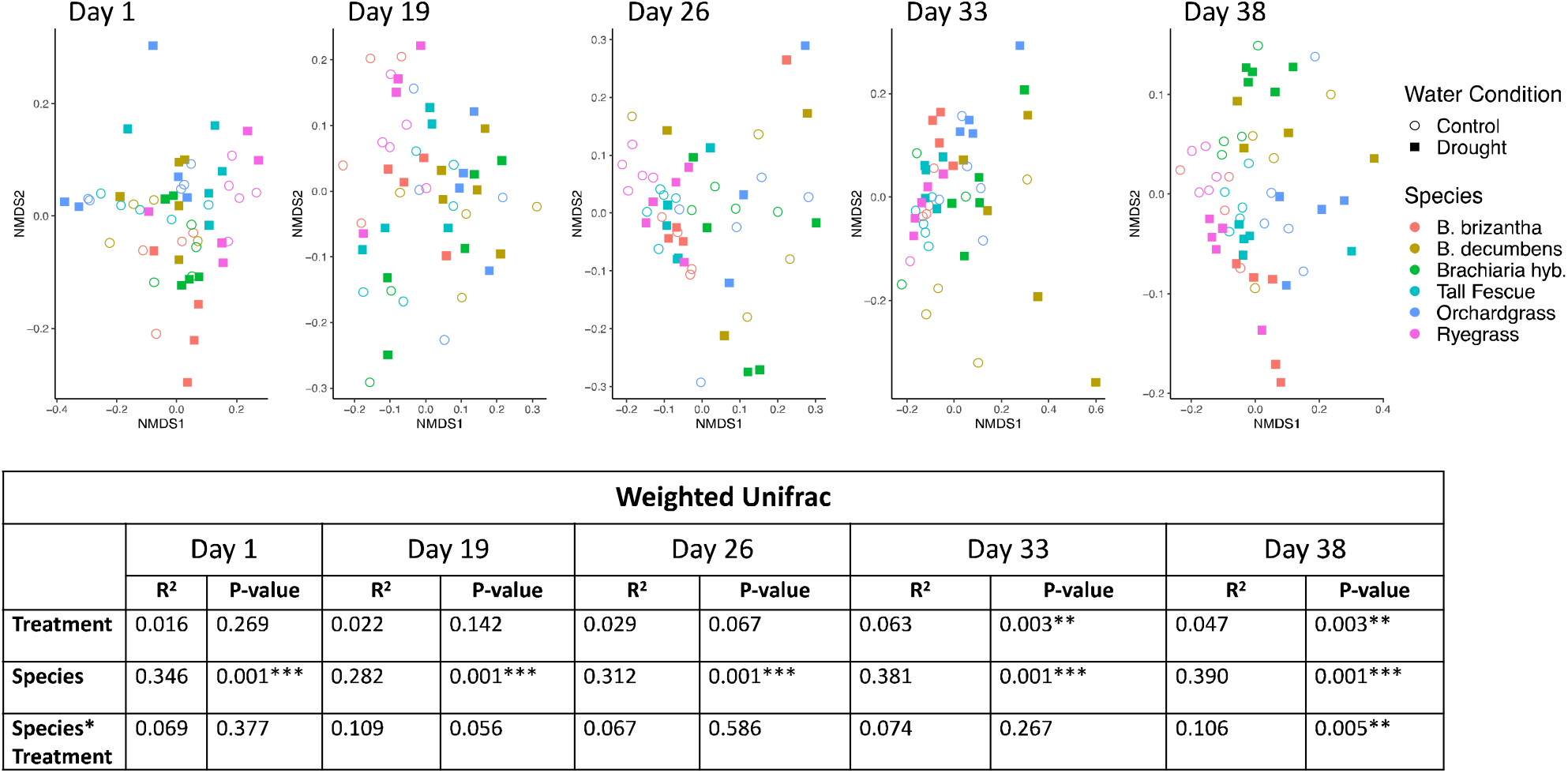
Bacterial communities from each host species became more distinct over time and were significantly impacted by drought stress. NMDS ordination was plotted for each sampling day using weighted UniFrac distances. PERMANOVA was conducted for each corresponding day to determine how communities were changing over time and when drought stress altered community structure.

To further understand changes in community structure, average community distance over time was modelled. On temperate grasses, average distance significantly decreased over time in both control and drought treatments (p.adj<0.001, TukeyHSD posthoc analyses). However, community distance on tropical grasses remained stable over time. Additionally, significant differences were not observed between temperate and tropical control groups but were observed between drought samples (p.adj=0.04) (Figure 3).

**Figure 3.**
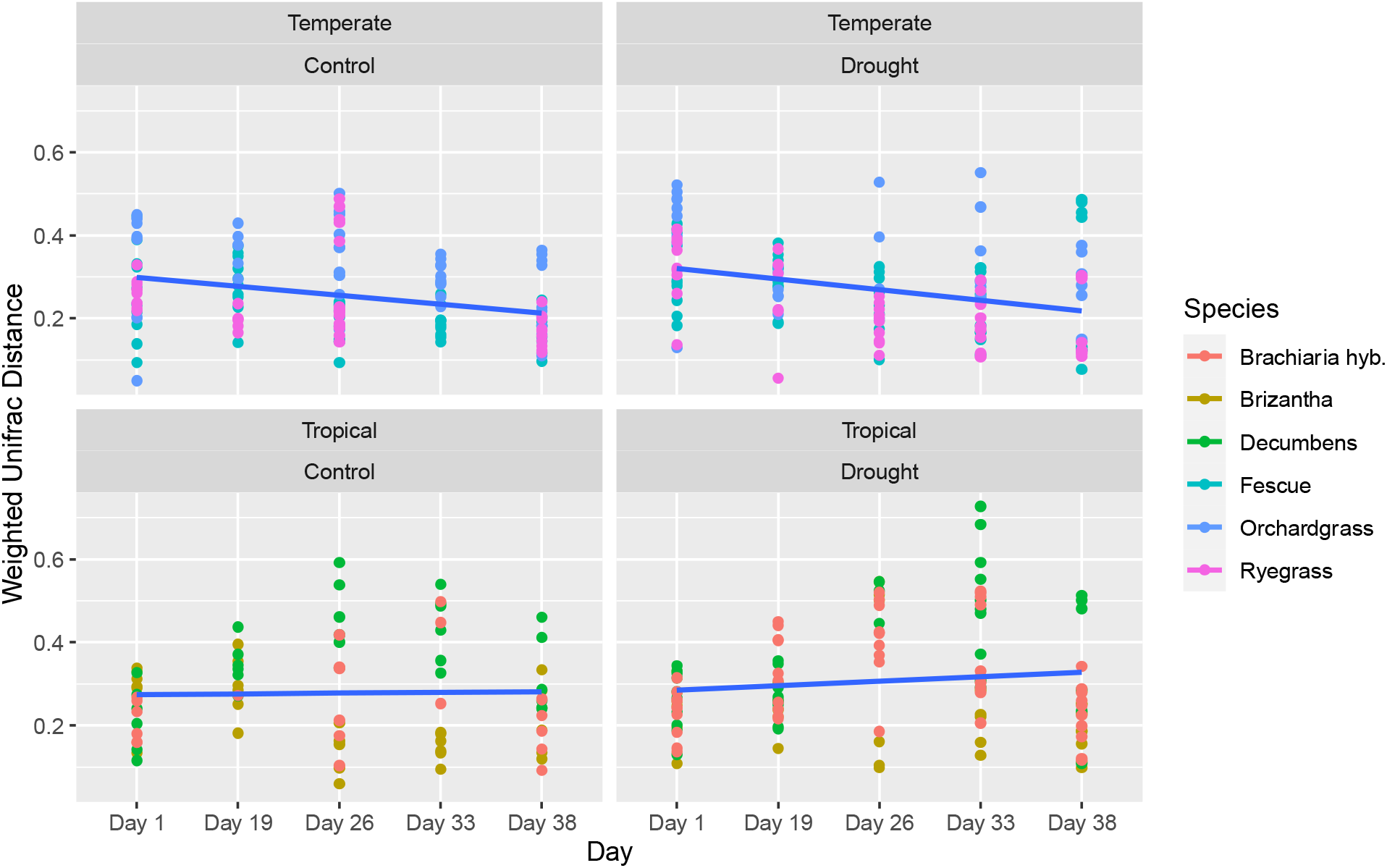
Phyllosphere communities became more similar over time regardless of treatment on the temperate host species but stayed the same on the tropical host species. Average distance between samples from the same host species were calculated for each sampling day using weighted UniFrac distance. Average distance within each host species significantly decreased in the communities from temperate hosts but did not change in the communities from the tropical hosts.

We analyzed individual host species to understand how susceptible to drought each bacterial community was based on host species. Ryegrass microbial communities were the first to show changes between control and drought treatments; significant differences were first observed on day 26 (R^2^= 0.37, P=0.012) (Supplementary Figure S3). *B. brizantha* (R^2^= 0.47, P=0.031), Tall Fescue (R^2^= 0.27, P=0.035), and Orchardgrass (R^2^=0.25, P=0.033) first showed significant differences on day 33; *Brachiaria* hybrid (R^2^= 0.30, P=0.006) on day 38; and *B. decumbens* never displayed significant differences.

### Machine Learning Allows Accurate Prediction of Microbial Communities in Drought

Machine learning (ML) allows detection of trends missed by traditional methods such as PERMANOVA [43], and allows identification of features that enable its predicative power. We used ML to test for a common response to drought among host species despite plant host selection on microbial communities. ML revealed high predictive power in determining if microbial communities were from the control or drought treatment (AUC=0.87) (Supplementary Figure S4) and that the top eight ASVs contributed 0.07 to our AUC value (Figure 4, Supplementary Table S2).

**Figure 4.**
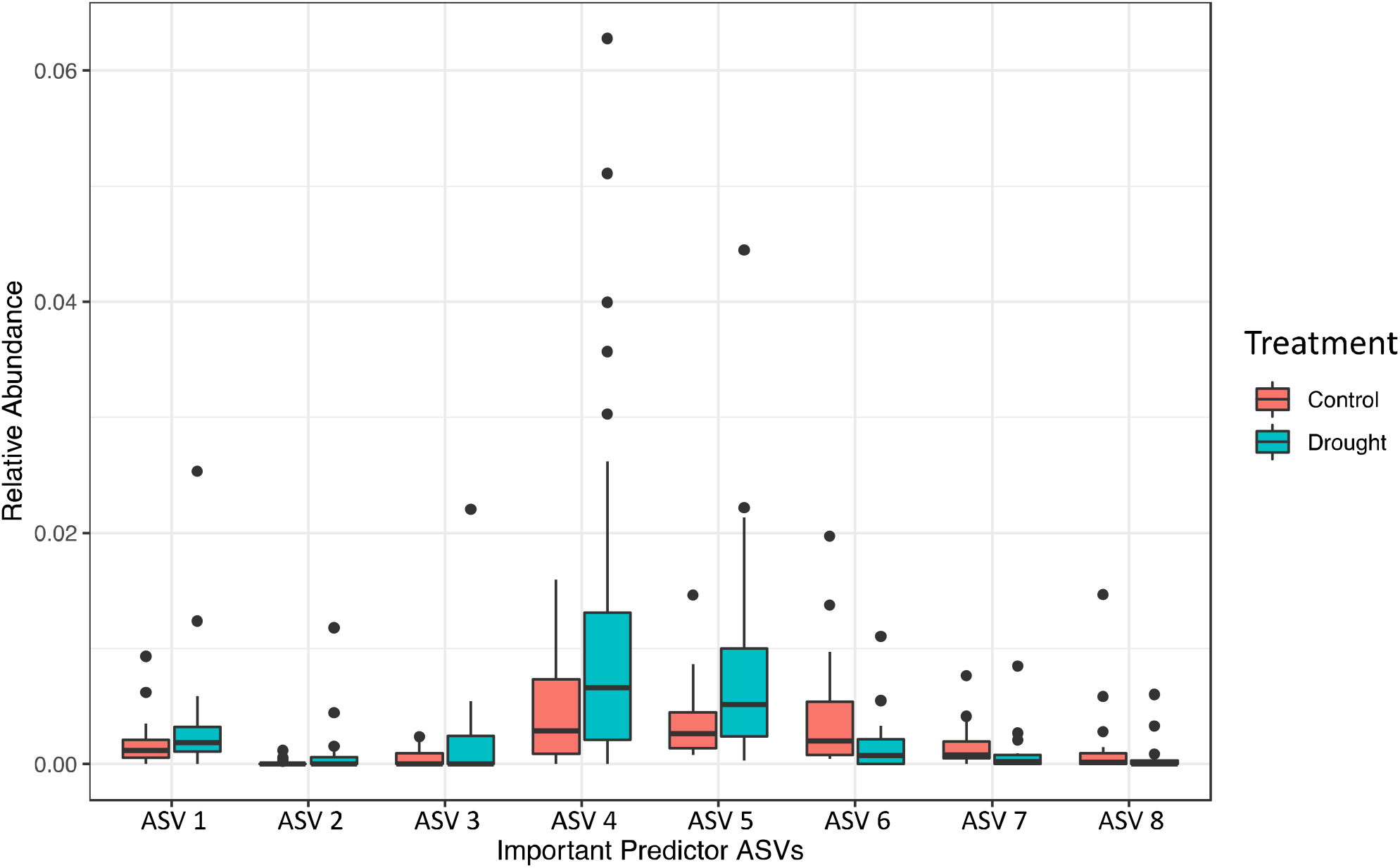
The top eight ASVs important for predicting if samples were from control or drought stressed plant hosts. Average relative abundance of each of the eight ASVs is given for control (n=27) and drought stressed plants (n=30) on day 38 of the experiment. ASV identities provided in Supplementary Table S1.

Additionally, ML had high predictive power in determining if communities were from temperate or tropical hosts (AUC=0.89) at the end of the experiment regardless of drought treatment. The model identified that 2 features, *Sphingomonas mali* and *Methylobacterium organophilum*, contributed 0.107 to our AUC values, indicating their presence was important in model performance (Supplementary Figure S5). Since tropical grasses are more related to each other than to the temperate grasses, this analysis helped determine if community assembly is stochastic or deterministic and identifies features associated with the two grass types.

### Grass Host Phylogeny Influences Phyllosphere Communities

Host species impact on community assembly and response to drought was tested using phylosymbiosis, which occurs when significant association between host species phylogeny and associated microbial communities occur [23]. Mantel tests on Bray-Curtis dissimilarities showed more closely related host species had more similar microbial communities (Mantel r=0.117, p=0.0001). Additionally, microbial communities were more related to host phylogeny during drought stress (Mantel r=0.202, p=0.007) than under control conditions (Mantel r=0.158, p=0.02) at the end of the drought period. Tests of phylosymbiosis using Weighted UniFrac measures showed similar trends but with weaker associations (All: Mantel r=0.064, p=0.0002; Drought: Mantel r=0.114, p=0.05; Control: Mantel r=0.057, p=0.19). Because weighted UniFrac incorporates phylogenetic information, it reduces nuanced variations at the tips of the bacterial phylogenetic trees [25].

To further explore evolutionary relationships between host phylogeny and bacterial communities, cophylogeny was tested with two separate global-fit methods. Global-fit methods test congruence between host phylogenetic trees and the corresponding microbial phylogeny and allow for identification of significant associations. PACo (Procrustes Approach to Cophylogeny) uses Procrustes analyses to test the dependency of one phylogeny on the other [29, 30]. ParaFit compares two distance matrices constructed from host and microbial phylogenetic distances and tests for random associations between the groups [52]. Positive correlations can indicate host-microbe coevolution [32, 53]. Tests for cophylogeny conducted on all samples collected on day 38 regardless of treatment using ParaFit (ParaFitGlobal=1.6024, p=0.001, permutations=999), and PACo (PACo=0.999, p=0.003) revealed significant global-fit cophylogenetic relationships.

The influence of drought stress on cophylogenetic signal was determined to understand if microbial response to drought was a stochastic process, and to look for evidence of a joint plant-microbial response. Results using Parafit from both control (ParaFitGlobal=1.136, p=0.001) and drought (ParaFitGlobal=1.296, p=0.001) showed evidence of cophylogeny. In the control treatment there were 414 significant associations between bacteria and plant hosts and 340 significant associations in drought treatment samples. Tanglegrams displaying significant associations between host and microbe phylogenies were created for control and drought treatments (Figure 5). Evidence of cophylogeny at the end of the experimental period was additionally detected using PACo for control (PACo=0.998, p=0.001) and drought (PACo=0.999, p=0.002) treatments.

**Figure 5.**
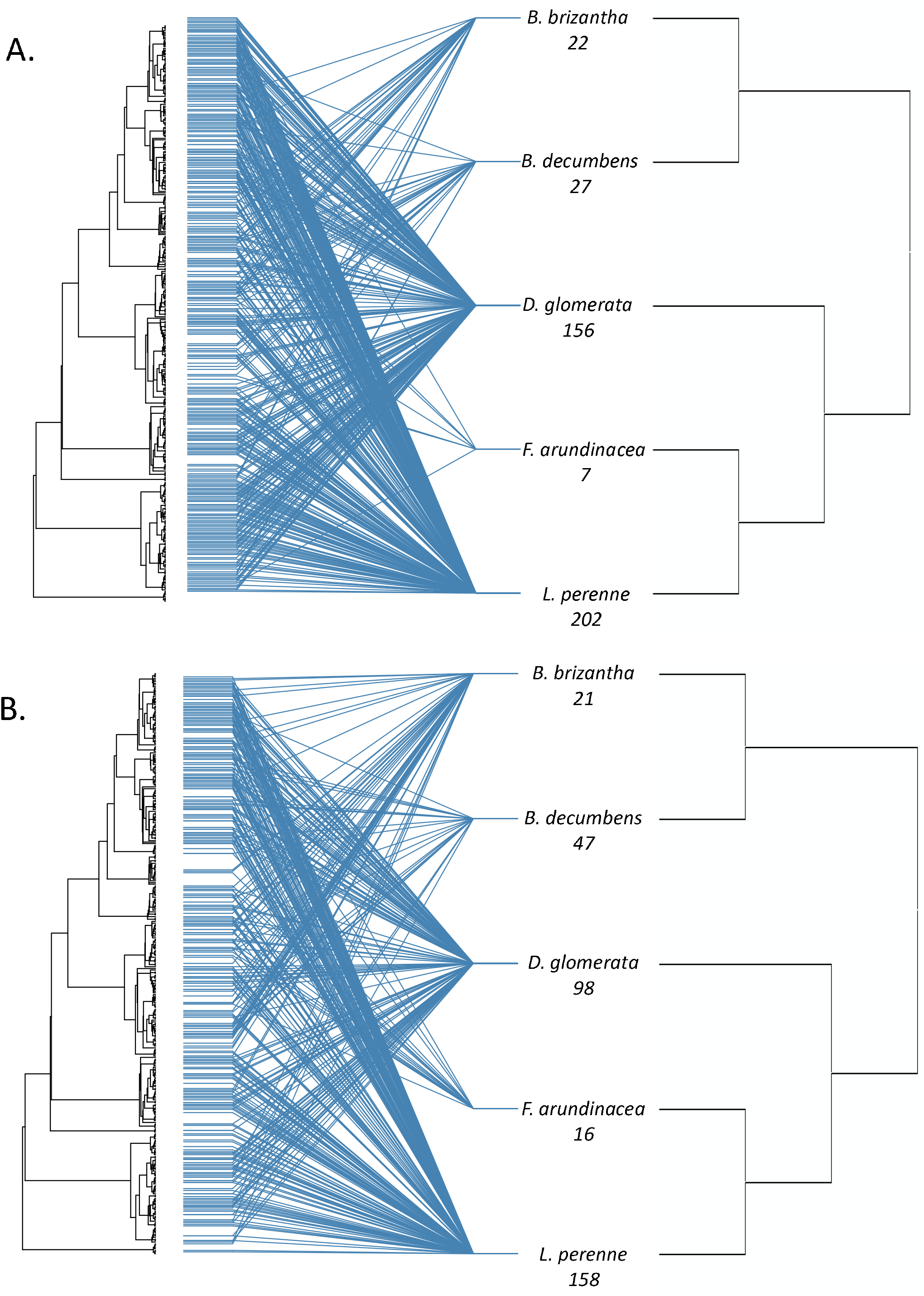
Cophylogenetic relationship analysis was conducted for (A) control plants (n=27) and (B) drought stressed plants (n=30). Blue lines in this tanglegram represent significant associations between phyllosphere bacteria on the left and their plant hosts on the right measured using ParaFitGlobal, which were determined if either of the ParaFit F statistics were below 0.05. Numbers under host species identity indicate the number of significant associations that a host species has with the bacterial phylogenetic tree. The bacterial phylogenetic tree was constructed in QIIME2 using FastTree which infers approximately-maximum-likelihood phylogenetic trees. The maximum-likelihood tree for the grass host phylogeny was constructed in MEGA. Only five grass species were included because host sequence information was not available for the *Brachiaria* hybrid.

### nifH gene abundance varies over time and by host species

No significant trend in *nifH* gene abundance was observed as a result of drought treatment, but significant differences in abundance were observed between host species and over time (Figure 6). The temperate grasses displayed a decrease in abundance over time to varying degrees, but the tropical grasses did not. Ryegrass control samples were temporally stable, but drought samples significantly decreased between day 1 and day 38 (p.adj=0.004). Tall Fescue (p.adj=0.02) and Orchardgrass (p.adj=0.006) significantly decreased between sample day 1 and 38 regardless of treatment. Control samples of *B. brizantha* showed no significant changes over time, but *nifH* copy number significantly increased over time in drought samples (p.adj=0.001). *Brachiaria* hyb. showed no differences as a result of drought but significantly varied across days (Day1 compared to 26 (p.adj=0.001) and day 33 (p.adj=0.03)). *B. decumbens* significantly increased over time in both control and drought conditions (p.adj>0.001). The trends over time for *nifH* abundance closely matched the trends observed in average UniFrac distance over time for each host species (Supplementary Figure S6).

**Figure 6.**
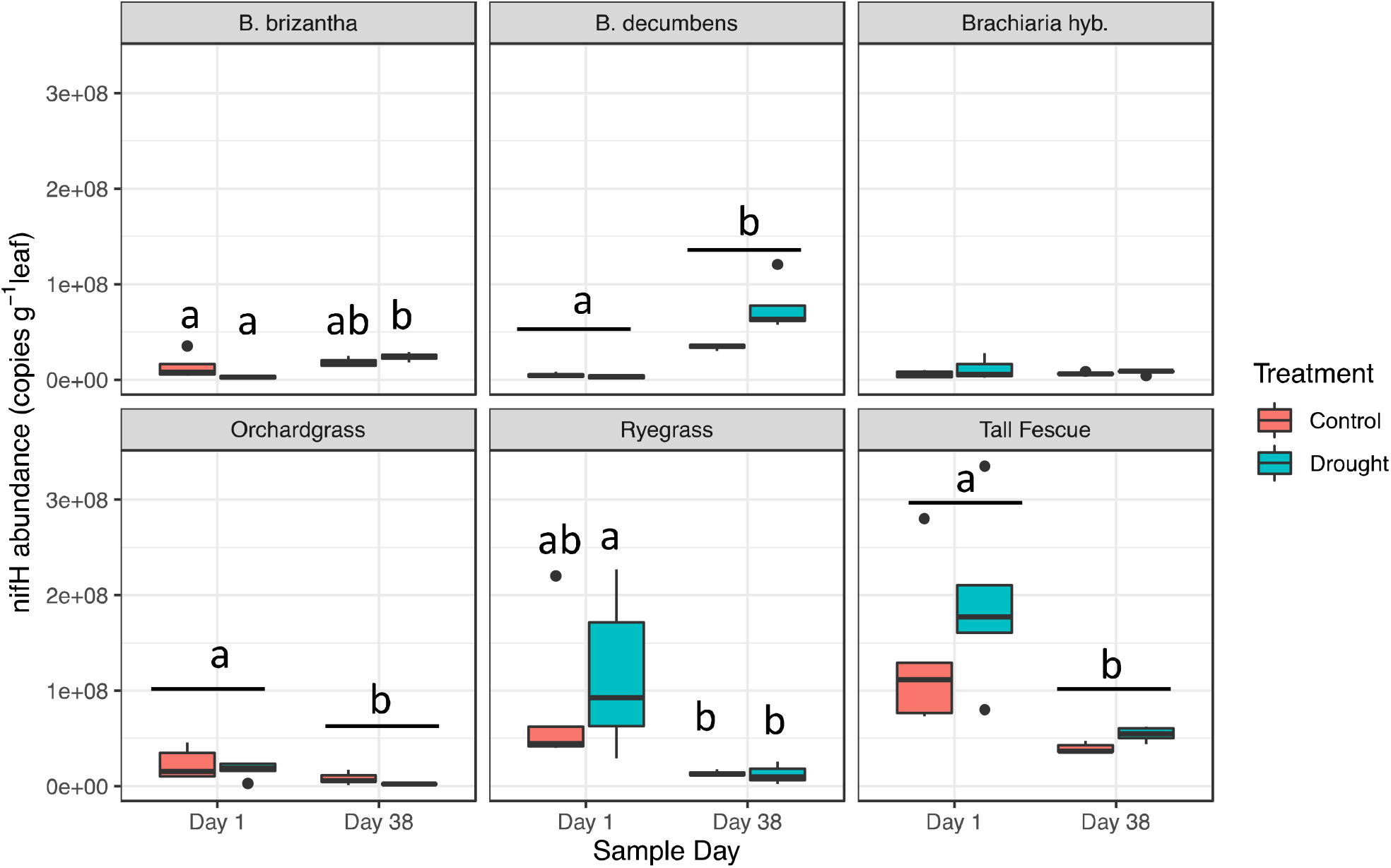
Abundance of the *nifH* gene was significantly different between host species and changed over time. However, it was not significantly impacted by drought stress. *nifH* abundance was measured using qPCR and standardized to number of copies per gram of leaf material.

## DISCUSSION

Evolutionary history impacts grass phyllosphere communities and their response to drought. Consistent with previous studies across multiple plant species, we found that host species was the most important factor influencing community assembly and that *Alphaproteobacteria* dominated communities [10, 13, 54, 55]. Additionally, our study revealed that communities changed over time and as a result of drought. We observed strong temporal patterns in which *Gammaproteobacteria* were replaced over time by *Cytophagia*, similar to studies in switchgrass that found *Gammaproteobacteria* were replaced throughout the growing season by *Alphaproteobacteria* [56]. While temporal replacement occurred on each host species, degree of replacement varied widely and between treatments. By the end of the experimental period, *Cytophagia* was the dominant class on control plants while *Alphaproteobacteria* dominated drought plants. The persistent presence of *Alphaproteobacteria*, in particular *Sphingomonas* and *Methylobacterium*, throughout the experiment on control and drought stressed plants likely resulted from niche partitioning and their complementary metabolisms uniquely suited to the phyllosphere [11, 56]. In the phyllosphere, *Sphingomonas* survive on a wide range of substrates due to high abundance of TonB receptors, while *Methylobacterium* can grow on one-carbon compounds such as methanol, a byproduct of host cell-wall metabolism [11, 57]. Additionally, their flexible metabolisms allow for adaptation to changing nutrient availability as leaf conditions change. Not only are they able to survive the harsh phyllosphere environment, they can promote plant growth and stress tolerance. Inoculation of *Sphingomonas* onto soybean plants resulted in increased tolerance of drought conditions and *Methylobacterium* on leaf surfaces are able to fix nitrogen and increase plant biomass production [58, 59]. The observed persistence under stress conditions in combination with their functional benefits, could indicate coevolutionary adaptation to life in the phyllosphere. Furthermore, *Sphingomonas* and *Methylobacterium* should be explored as biofertilizers because of their widespread presence and observed drought tolerance.

Host species effect on community assembly increased over time. On day one of the experiment, host species accounted for 38% of community variability in the control samples compared to 57% on the last day. This likely results from host selection on community assembly; host species selection increases over time as communities successfully establish, as more bacteria land on the leaf surface through dispersal, and as communities change in relation to plant development [56, 60, 61]. Interestingly, the effect of host species overtime was different between the native temperate grasses and the non-native tropical grasses. On the first day of the experiment, host species exhibited similar influence on microbial communities from temperate and tropical grasses. However, by the end of the experimental period, species explained 72% of the variability on temperate grass hosts but only 39% on tropical grass hosts. The difference in effect over time between the tropical and temperate grasses likely results from host-microbe evolutionary relationships that exist for the native temperate species but not for the non-native tropical species.

To understand if phyllosphere communities from temperate grasses experienced increased selection compared to tropical grasses, we determined how ecological distance changed over time for each host species. Since temperate grasses were grown in their native environment, we expected increased host selection compared to tropical grasses. While change over time accounted for similar amounts of variability in the tropical (27%) and temperate grasses (23%), the average distance of communities found on each host species decreased in temperate grasses but remained stable in tropical grasses. These significant differences suggest deterministic assembly in the phyllosphere. In the temperate grasses, decreased distance could result from increased selection caused by coevolved plant-microbe relationships. However, since tropical grasses were not grown in an environment with their native microbiota, changes over time and between species were more likely a result of host physiology.

Presence of phylosymbiosis under non-stressed conditions indicates host-species influences community assembly and that bacterial communities are more similar to each other on plant hosts that are more phylogenetically similar [62]. While phylosymbiosis could result from coevolution or cospeciation, it can also result from differences in host ecological niche, geographic locations, or host filtering in which related hosts have many shared traits [23, 63, 64]. Therefore, phylosymbiosis does not determine a specific mechanism. Presence of phylosymbiosis demonstrates that phyllosphere community assembly is a deterministic process but what is driving it is still not fully understood. By growing plants in the same environment, we eliminated some of the confounding factors that might otherwise contribute to this relationship such as differences in soil, weather patterns, or biogeographic separation. Previous work across animal species concluded that when related hosts grown under identical conditions maintain distinct microbial communities, it is analogous to microbial markers of host evolutionary relationships [26].

Unsurprisingly, the cophylogenetic analysis revealed strong evidence of cophylogeny in the temperate grass species with hundreds of significant correlations compared to only dozens observed in the tropical grasses. Thus, cophylogenetic signal was stronger in the native temperate grasses than in non-native tropical grasses. The differences between tropical and temperate hosts further supports the idea that the host-species effect seen in the temperate grasses is a result of coevolution as microbial members have adapted alongside their host.

### Microbial Community Response: An Adaptation to Drought

Understanding how phyllosphere communities respond to drought in relation to their plant host is important for understanding how we can use bacteria as biofertilizers to promote plant health in the future. Interestingly, no difference in alpha diversity as a result of drought was observed even though drought caused changes in bacterial community structures. Previous work found that phyllosphere community diversity but not composition was related to plant community productivity [6]. Therefore, shifts we are seeing in community structure but not in alpha diversity could indicate microbial communities act as a stress response trait.

The forage grass species used in this experiment have varying degrees of drought tolerance resulting from the different strategies used under drought stress. Drought tolerance is well documented for the temperate grasses used in this study. Ryegrass is the most drought susceptible and under field conditions tall fescue is the most drought tolerant [65, 66]. However, when grown in pots, orchardgrass exhibits higher drought tolerance due to its abilities to take up water in low soil moisture conditions, promote membrane stabilization, and protect its meristem from dehydration [66, 67]. In the field, tall fescue has higher drought tolerance due to its ability to form deep root networks, which were limited by the depth of pots in which they were grown. The C4 tropical plants have greater water use efficiency due to their ability to maintain higher photosynthetic rates under decreased water stress compared to the C3 temperate species [68, 69]. Additionally, they can form extensive root networks that enable high water uptake efficiency from soil [70]. These levels of known drought tolerance correlate with the changes we saw in microbial community structures. Ryegrass, the most drought susceptible, was the first to show signs of change, followed by orchardgrass and tall fescue. The more drought tolerant tropical grasses showed changes in microbial communities later than the temperate grasses. Even though previous studies found *B. brizantha* and *B. decumbens* to have similar drought tolerances, we observed changes in *B. brizantha* communities on day 33 but no significant changes as a result of drought in *B. decumbens* [70, 71]. What remains unclear is, if changes in microbial community structure are in direct response to drought or in response to changes in host physiology.

If changes in communities were in direct response to drought, we may expect to see a decrease in phylosymbiosis as selection imposed by host species decreased and communities became more similar to each other. Instead, we observed an increase in phylosymbiosis under drought stress indicating that selection on microbial communities is increasing. Conversely, total cophylogenetic associations decreased as a result of drought with less overall connections between host and microbe phylogenies. However, not all host species showed similar trends, indicating that microbial community response to drought, much like community assembly, is a deterministic process facilitated by changes in plant host physiology. Because strong evidence of phylosymbiosis and cophylogeny remain in spite of shifts in community structure, we propose that phyllosphere communities are a plant stress response trait that has coevolved alongside its plant host.

Despite the strong host species effect on microbial communities during drought, we used machine learning to determine if there was a common response to drought across our host species. Determining if any microbes invariably survive under drought conditions across our range of hosts and have the potential to promote plant growth is important for determining prospective bacteria to test as biofertilizers. Our ML pipeline accurately predicted if a sample came from a drought stressed or control plant 87% of the time, confirming that there is a common response to drought despite divergent communities. No single bacterial ASV was responsible for model prediction, rather several ASVs provided minor predictive power. Of the top 8 predictors, 5 from the order Actinomycetales were slightly elevated in drought samples including *Microbacterium*. In the rhizosphere, *Microbacterium* can produce volatile compounds that promote plant health and growth [72] and help regulate plant response to drought stress by altering the metabolite profile to promote osmoregulation [73]. Thus, even though microbial communities are host specific, core functions exist in phyllosphere communities across plant hosts that enable microbial survival under harsh conditions while also offering functional support to their plant host.

### Nitrogen Fixation: A Core Function

We used nitrogen fixation as one example of an important function microbes provide plants. Recent studies found phyllosphere communities input nitrogen into their ecosystems [9, 59, 74]. When we assessed our communities for nitrogen fixation potential, we found stable diazotroph presence on every host species. However, *nifH* abundance was not negatively impacted by drought. The occurrence of temporally stable diazotrophs on every host species indicates its likely an important part of the functional core community, while the differential abundance between host species points to the evolutionary relationships between plants and their microbes.

When under drought stress, the most drought tolerant host species, *B. decumbens*, exhibited increased *nifH* abundance and a strong cophylogenetic relationship with the bacterial family Oxalobacteraceae accounting for 14 of the 47 significant relationships. Additionally, relative abundance of Oxalobacteraceae from the class Betaproteobacteria increased on *B. decumbens* under drought. Oxalobacteraceae are adapted to oligotrophic conditions and some genera are nitrogen-fixers (Supplementary Figure S7) [75]. Because of the sustained presence of *nifH* across time and treatments in combination with their correlation to community structure, we propose nitrogen-fixation as a keystone function of phyllosphere communities. This further supports our hypothesis that microbial communities are a plant trait which help promote plant stress tolerance.

## Conclusion

This study revealed phyllosphere community assembly is related to host evolutionary history. The strong evidence of phylosymbiosis in combination with increased selection and cophylogeny in the native temperate grasses compared to the non-native tropical grasses, suggests that microbial communities are a plant trait that coevolve alongside their plant hosts. The conserved presence but differential abundance of important bacteria such as *Sphingomonas* and *Methylobacterium*, and the functional potential of nitrogen fixation during drought stress further support the idea that microbial communities are plant traits that evolve to promote plant growth and stress tolerance. Future studies should look at the effect of inoculating plants with the taxonomically and functionally important bacteria identified in this study that were also temporally and drought stable. Creating biofertilizers with ecologically important and evolutionarily selected microbes could promote plant health and tolerance to a changing climate.

## ACKNOWLEDGMENTS

We would like to thank Michelle DaCosta and Jefferson Lu for guidance and equipment for measuring plant health. Genomics service and sequencing was performed by Ravi Ranjan at the Genomics Resource Laboratory, University of Massachusetts Amherst, MA. We are thankful for financial support from the Lotta M. Crabtree Foundation, and the National Science Foundation – Dimensions of Biodiversity (DEB 1442183).

## SUPPLEMENTARY INFORMATION

Supplementary information is available online only.

Single PDF file, 5.1 Mb of Supplementary information containing table of contents, methods, tables, figures, and references.

## REFERENCES

1. Bengtsson J, Bullock JM, Egoh B, Everson C, Everson T, O’Connor T, et al. Grasslands— more important for ecosystem services than you might think. Ecosphere 2019; 10: e02582.

2. Ciais Ph, Reichstein M, Viovy N, Granier A, Ogée J, Allard V, et al. Europe-wide reduction in primary productivity caused by the heat and drought in 2003. Nature 2005; 437: 529–533.

3. Steiger NJ, Smerdon JE, Cook BI, Seager R, Williams AP, Cook ER. Oceanic and radiative forcing of medieval megadroughts in the American Southwest. Science Advances 2019; 5: eaax0087.

4. Wolfe DW, DeGaetano AT, Peck GM, Carey M, Ziska LH, Lea-Cox J, et al. Unique challenges and opportunities for northeastern US crop production in a changing climate. Climatic Change 2018; 146: 231–245.

5. Smith MD. The ecological role of climate extremes: current understanding and future prospects. Journal of Ecology 2011; 99: 651–655.

6. Laforest-Lapointe I, Paquette A, Messier C, Kembel SW. Leaf bacterial diversity mediates plant diversity and ecosystem function relationships. Nature 2017; 546: 145–147.

7. Saleem M, Meckes N, Pervaiz ZH, Traw MB. Microbial Interactions in the Phyllosphere Increase Plant Performance under Herbivore Biotic Stress. Front Microbiol 2017; 8.

8. Rastogi G, Coaker GL, Leveau JHJ. New insights into the structure and function of phyllosphere microbiota through high-throughput molecular approaches. FEMS Microbiology Letters 2013; 348: 1–10.

9. Fürnkranz M, Wanek W, Richter A, Abell G, Rasche F, Sessitsch A. Nitrogen fixation by phyllosphere bacteria associated with higher plants and their colonizing epiphytes of a tropical lowland rainforest of Costa Rica. ISME J 2008; 2: 561–570.

10. Redford AJ, Bowers RM, Knight R, Linhart Y, Fierer N. The ecology of the phyllosphere: geographic and phylogenetic variability in the distribution of bacteria on tree leaves. Environ Microbiol 2010; 12: 2885–2893.

11. Delmotte N, Knief C, Chaffron S, Innerebner G, Roschitzki B, Schlapbach R, et al. Community proteogenomics reveals insights into the physiology of phyllosphere bacteria. Proceedings of the National Academy of Sciences 2009; 106: 16428–16433.

12. Aydogan EL, Moser G, Müller C, Kämpfer P, Glaeser SP. Long-Term Warming Shifts the Composition of Bacterial Communities in the Phyllosphere of Galium album in a Permanent Grassland Field-Experiment. Front Microbiol 2018; 9.

13. Kembel SW, O’Connor TK, Arnold HK, Hubbell SP, Wright SJ, Green JL. Relationships between phyllosphere bacterial communities and plant functional traits in a neotropical forest. PNAS 2014; 111: 13715–13720.

14. Lajoie G, Maglione R, Kembel SW. Adaptive matching between phyllosphere bacteria and their tree hosts in a neotropical forest. Microbiome 2020; 8: 70.

15. Rosenberg E, Zilber-Rosenberg I. The hologenome concept of evolution after 10 years. Microbiome 2018; 6: 78.

16. Lindow SE, Brandl MT. Microbiology of the Phyllosphere. Appl Environ Microbiol 2003; 69: 1875–1883.

17. Lindow SE, Leveau JHJ. Phyllosphere microbiology. Current Opinion in Biotechnology 2002; 13: 238–243.

18. Rosado BHP, Almeida LC, Alves LF, Lambais MR, Oliveira RS. The importance of phyllosphere on plant functional ecology: a phyllo trait manifesto. New Phytologist 2018; 219: 1145–1149.

19. Li Y, Sun H, Wu Z, Li H, Sun Q. Urban traffic changes the biodiversity, abundance, and activity of phyllospheric nitrogen-fixing bacteria. Environ Sci Pollut Res 2019; 26: 16097–16104.

20. Atamna-Ismaeel N, Finkel O, Glaser F, Mering C von, Vorholt JA, Koblížek M, et al. Bacterial anoxygenic photosynthesis on plant leaf surfaces. Environmental Microbiology Reports 2012; 4: 209–216.

21. Carvalho SD, Castillo JA. Influence of Light on Plant–Phyllosphere Interaction. Front Plant Sci 2018; 9.

22. Vandenkoornhuyse P, Quaiser A, Duhamel M, Van AL, Dufresne A. The importance of the microbiome of the plant holobiont. New Phytologist 2015; 206: 1196–1206.

23. Lim SJ, Bordenstein SR. An introduction to phylosymbiosis. Proc R Soc B 2020; 287: 20192900.

24. Brucker RM, Bordenstein SR. The Hologenomic Basis of Speciation: Gut Bacteria Cause Hybrid Lethality in the Genus Nasonia. Science 2013; 341: 667–669.

25. Pollock FJ, McMinds R, Smith S, Bourne DG, Willis BL, Medina M, et al. Coral-associated bacteria demonstrate phylosymbiosis and cophylogeny. Nat Commun 2018; 9: 4921.

26. Brooks AW, Kohl KD, Brucker RM, Opstal EJ van, Bordenstein SR. Phylosymbiosis: Relationships and Functional Effects of Microbial Communities across Host Evolutionary History. PLOS Biology 2016; 14: e2000225.

27. Moran NA, Sloan DB. The Hologenome Concept: Helpful or Hollow? PLOS Biology 2015; 13: e1002311.

28. Page RDM. Tangled Trees: Phylogeny, Cospeciation, and Coevolution. 2003. University of Chicago Press.

29. Hutchinson MC, Cagua EF, Balbuena JA, Stouffer DB, Poisot T. paco: implementing Procrustean Approach to Cophylogeny in R. Methods in Ecology and Evolution 2017; 8: 932–940.

30. Balbuena JA, Míguez-Lozano R, Blasco-Costa I. PACo: A Novel Procrustes Application to Cophylogenetic Analysis. PLOS ONE 2013; 8: e61048.

31. Weckstein JD. Biogeography Explains Cophylogenetic Patterns in Toucan Chewing Lice. Systematic Biology 2004; 53: 154–164.

32. Youngblut ND, Reischer GH, Walters W, Schuster N, Walzer C, Stalder G, et al. Host diet and evolutionary history explain different aspects of gut microbiome diversity among vertebrate clades. Nat Commun 2019; 10.

33. Innerebner G, Knief C, Vorholt JA. Protection of Arabidopsis thaliana against Leaf-Pathogenic Pseudomonas syringae by Sphingomonas Strains in a Controlled Model System. Appl Environ Microbiol 2011; 77: 3202–3210.

34. Noble AS, Noe S, Clearwater MJ, Lee CK. A core phyllosphere microbiome exists across distant populations of a tree species indigenous to New Zealand. PLoS One 2020; 15.

35. Barrs HD, Weatherley PE. A Re-Examination of the Relative Turgidity Technique for Estimating Water Deficits in Leaves. Aust Jnl Of Bio Sci 1962; 15: 413–428.

36. Hoffman L, DaCosta M, Ebdon JS, Watkins E. Physiological Changes during Cold Acclimation of Perennial Ryegrass Accessions Differing in Freeze Tolerance. Crop Science 2010; 50: 1037–1047.

37. Lutts S, Kinet JM, Bouharmont J. NaCl-induced Senescence in Leaves of Rice (Oryza sativa L.) Cultivars Differing in Salinity Resistance. Annals of Botany 1996; 78: 389–398.

38. Naqib A, Poggi S, Wang W, Hyde M, Kunstman K, Green SJ. Making and Sequencing Heavily Multiplexed, High-Throughput 16S Ribosomal RNA Gene Amplicon Libraries Using a Flexible, Two-Stage PCR Protocol. Methods Mol Biol 2018; 1783: 149–169.

39. Poly F, Monrozier LJ, Bally R. Improvement in the RFLP procedure for studying the diversity of nifH genes in communities of nitrogen fixers in soil. Research in Microbiology 2001; 152: 95–103.

40. Bolyen E, Rideout JR, Dillon MR, Bokulich NA, Abnet CC, Al-Ghalith GA, et al. Reproducible, interactive, scalable and extensible microbiome data science using QIIME 2. Nature Biotechnology 2019; 37: 852–857.

41. Topçuoğlu BD, Lapp Z, Kelly Sovacool, Evan Snitkin, Jenna Wiens, Patrick D. Schloss. mikRopML: User-Friendly R Package for Robust Machine Learning Pipelines. Journal of Open Source Software 2021; 6: 3073.

42. R Core Team. R: a language and environment for statistical computing. https://www.R-project.org/. Accessed 31 Mar 2020.

43. Topçuog BD. A Framework for Effective Application of Machine Learning to Microbiome-Based Classification Problems. 2020; 11: 13.

44. Stecher G, Tamura K, Kumar S. Molecular Evolutionary Genetics Analysis (MEGA) for macOS. Molecular Biology and Evolution 2020; 37: 1237–1239.

45. Schoch CL, Ciufo S, Domrachev M, Hotton CL, Kannan S, Khovanskaya R, et al. NCBI Taxonomy: a comprehensive update on curation, resources and tools. Database (Oxford) 2020; 2020.

46. Edgar RC. MUSCLE: multiple sequence alignment with high accuracy and high throughput. Nucleic Acids Res 2004; 32: 1792–1797.

47. Oksanen J, Blanchet FG, Friendly M, Kindt R, Legendre P, McGlinn D, et al. vegan: Community Ecology Package. 2019.

48. Paradis E, Schliep K. ape 5.0: an environment for modern phylogenetics and evolutionary analyses in R. Bioinformatics 2019; 35: 526–528.

49. Bates D, Mächler M, Bolker B, Walker S. Fitting Linear Mixed-Effects Models Using lme4. Journal of Statistical Software 2015; 67: 1–48.

50. Lenth RV. Least-Squares Means: The R Package lsmeans. Journal of Statistical Software 2016; 69: 1–33.

51. Wickham H. ggplot2: Elegant Graphics for Data Analysis. 2016. Springer-Verlag New York.

52. Legendre P, Desdevises Y, Bazin E. A Statistical Test for Host–Parasite Coevolution. Systematic Biology 2002; 51: 217–234.

53. Abdullaeva Y, Ambika Manirajan B, Honermeier B, Schnell S, Cardinale M. Domestication affects the composition, diversity, and co-occurrence of the cereal seed microbiota. Journal of Advanced Research 2020.

54. Laforest-Lapointe I, Messier C, Kembel SW. Tree Leaf Bacterial Community Structure and Diversity Differ along a Gradient of Urban Intensity. mSystems 2017; 2.

55. Kim M, Singh D, Lai-Hoe A, Go R, Abdul Rahim R, A.N. A, et al. Distinctive Phyllosphere Bacterial Communities in Tropical Trees. Microb Ecol 2012; 63: 674–681.

56. Grady KL, Sorensen JW, Stopnisek N, Guittar J, Shade A. Assembly and seasonality of core phyllosphere microbiota on perennial biofuel crops. Nat Commun 2019; 10: 4135.

57. Galbally IE, Kirstine W. The Production of Methanol by Flowering Plants and the Global Cycle of Methanol. Journal of Atmospheric Chemistry 2002; 43: 195–229.

58. Asaf S, Khan AL, Khan MA, Imran QM, Yun B-W, Lee I-J. Osmoprotective functions conferred to soybean plants via inoculation with Sphingomonas sp. LK11 and exogenous trehalose. Microbiological Research 2017; 205: 135–145.

59. Madhaiyan M, Alex THH, Ngoh ST, Prithiviraj B, Ji L. Leaf-residing Methylobacterium species fix nitrogen and promote biomass and seed production in Jatropha curcas. Biotechnol Biofuels 2015; 8.

60. Maignien L, DeForce EA, Chafee ME, Eren AM, Simmons SL. Ecological Succession and Stochastic Variation in the Assembly of Arabidopsis thaliana Phyllosphere Communities. mBio 2014; 5.

61. Ottesen AR, Gorham S, Reed E, Newell MJ, Ramachandran P, Canida T, et al. Using a Control to Better Understand Phyllosphere Microbiota. PLOS ONE 2016; 11: e0163482.

62. Brucker RM, Bordenstein SR. The roles of host evolutionary relationships (genus: Nasonia) and development in structuring microbial communities. Evolution 2012; 66: 349–362.

63. Groussin M, Mazel F, Alm EJ. Co-evolution and Co-speciation of Host-Gut Bacteria Systems. Cell Host & Microbe 2020; 28: 12–22.

64. Mazel F, Davis KM, Loudon A, Kwong WK, Groussin M, Parfrey LW. Is Host Filtering the Main Driver of Phylosymbiosis across the Tree of Life? mSystems 2018; 3.

65. Cyriac D, Hofmann RW, Stewart A, Sathish P, Winefield CS, Moot DJ. Intraspecific differences in long-term drought tolerance in perennial ryegrass. PLoS One 2018; 13.

66. Volaire F, Lelièvre F. Drought survival in Dactylis glomerata and Festuca arundinacea under similar rooting conditions in tubes. Plant and Soil 2001; 229: 225–234.

67. Volaire F, Thomas H, Lelièvre F. Survival and recovery of perennial forage grasses under prolonged Mediterranean drought: I. Growth, death, water relations and solute content in herbage and stubble. New Phytologist 1998; 140: 439–449.

68. Taylor SH, Franks PJ, Hulme SP, Spriggs E, Christin PA, Edwards EJ, et al. Photosynthetic pathway and ecological adaptation explain stomatal trait diversity amongst grasses. New Phytologist 2012; 193: 387–396.

69. Cardoso JA, Pineda M, Jiménez J de la C, Vergara MF, Rao IM. Contrasting strategies to cope with drought conditions by two tropical forage C4 grasses. AoB Plants 2015; 7.

70. Guenni O, Marin DP, Baruch Z. Responses to drought of five Brachiaria species. I. Biomass production, leaf growth, root distribution, water use and forage quality. Plant and Soil 2002.

71. Guenni O, Baruch Z, Marín D. Responses to drought of five Brachiaria species. II. Water relations and leaf gas exchange. Plant and Soil 2004; 258: 249–260.

72. Cordovez V, Schop S, Hordijk K, Dupré de Boulois H, Coppens F, Hanssen I, et al. Priming of Plant Growth Promotion by Volatiles of Root-Associated Microbacterium spp. Appl Environ Microbiol 2018; 84.

73. Vílchez JI, Niehaus K, Dowling DN, González-López J, Manzanera M. Protection of Pepper Plants from Drought by Microbacterium sp. 3J1 by Modulation of the Plant’s Glutamine and α-ketoglutarate Content: A Comparative Metabolomics Approach. Front Microbiol 2018; 9.

74. Stanton DE, Batterman SA, Fischer JCV, Hedin LO. Rapid nitrogen fixation by canopy microbiome in tropical forest determined by both phosphorus and molybdenum. Ecology 2019; 100: e02795.

75. Baldani JI, Rouws L, Cruz LM, Olivares FL, Schmid M, Hartmann A. The Family Oxalobacteraceae. In: Rosenberg E, DeLong EF, Lory S, Stackebrandt E, Thompson F (eds). The Prokaryotes: Alphaproteobacteria and Betaproteobacteria. 2014. Springer, Berlin, Heidelberg, pp 919–974.

