## Supplemental Material for "Evolutionary History Impacts Phyllosphere Community Assembly on Forage Grasses"

**Contents:	Page**

Supplementary Methods	1-2

Supplementary Tables S1 – S2	3

Supplementary Figures S1 -S7	4-9

References	9

**Supplementary Methods**

*Transplanting Grasses*

In June 2019, all individual plants were transplanted into 15x30cm pots filled with soil collected from natural grass fields in Amherst, MA. The top 15 cm of topsoil was collected, all rocks and roots were removed, and soil was immediately placed in pots and allowed to settle for two days before plants were transplanted. Soil nutrient contents were tested at the University of Massachusetts Soil and Plant Nutrient Testing Laboratory (Supplementary Table S1). Before transplanting, roots and shoots were each trimmed to a uniform length of three inches. For the three temperate grass species, there were five replicates per plant species per treatment. For the three tropical grasses, four pots were used in the ‘control’ treatment and five were used in the ‘drought’ treatment. This resulted in 57 total pots.

*Library Preparation*

All phyllosphere community DNA samples underwent a two-step PCR amplification to attach barcodes and Illumina adaptor sequences. In the first PCR step, chloroplast excluding primers targeting the V5-V6 region of the 16S rRNA gene with linker sequences were used. The primer pair consisted of the forward primer (799F): 5’ACACTGACGACATGGTTCTACA AACMGGATTAGATACCCKG*-3’,* and the reverse primer (1115R): TACGGTAGCAGAGACTTGGTCT  AGGGTTGCGCTCGTTG*-3’*, where the underlined portion is the linker sequence followed by the primer sequence [1, 2]. The first PCR step was a 30 µl reaction containing 3 µl of 10X OmniKlentaq buffer (DNA Polymerase Technology, St. Louis, MO), 2.4 µl dNTPs (10 µM), 0.75 µl forward primer (5 µM), 0.75 µl reverse primer (5 µM), 0.5 µl bovine serum albumin (BSA), 0.2 µl OmniKlentaq LA, 3 µl DNA, and 19.4 µl molecular-grade water. The PCR reaction had an initial denaturation step at 95°C for 2 min followed by 30 cycles of 95°C for 20 s, 60°C for 30 s, 68°C for 30 s, and a final elongation step for 3 min at 68°C. The product from this PCR was used as the DNA template for the second PCR step, which attached individual barcodes to each sample to allow separation of samples in downstream analyses. The second PCR step was a 20 µl reaction containing 10 µl of 2X Dreamtaq Mastermix (Thermo Fisher Scientific, Waltham, MA), 4 µl water, 2 µl DNA, and 4 µl of Access Array Barcode primer pools (Fluidigm, San Francisco, CA). The second PCR reaction consisted of an initial 5 min denaturation step at 95°C; 8 cycles of 95°C for 30 s, 60°C for 30 s, and 72°C for 30 s; and a final 7 min elongation step at 72°C. The amplicons were pooled and sequenced on Illumina MiSeq Platform, with 251 bp paired-end sequencing chemistry. Illumina PhiX was spiked-in (~ 25%) to account for the low base diversity. Sequencing was performed at the Genomics Resource Laboratory (University of Massachusetts-Amherst).

*Quantitative PCR of Nitrogen Fixing Bacteria*

The abundance of nitrogen-fixing bacteria and the corresponding effect of drought stress were determined using qPCR quantifying the *nifH* gene. All samples were amplified in triplicate using the PolF(5′ TGC GAY CCS AAR GCB GAC TC 3′) and PolR primers(5′ ATS GCC ATC ATY TCR CCG GA 3′) in a 20 µl reaction containing 10 µl Luna qPCR mastermix (NEB, Ipswitch MA, USA), 1.5 µl forward primer (10 µm), 1.5 µl reverse primer (10 µm), 1 µl DNA, and 6 µl molecular-grade water. The qPCR reaction was carried out using 5 min of initial denaturation at 95°C, 40 cycles of 95°C for 30s, 60°C for 30s, and 72°C for 30s, followed by a final 10 min elongation step at 95°C. Abundance of *nifH* genes was normalized to an average mass of leaf samples used during DNA extraction for each host species.

**Supplementary Tables**

**Supplementary Table S1**. Soil physical-chemical analysis for experiments with tropical and temperate grasses. Macronutrient and micronutrient concentrations were determined using Modified Morgan extractables.

| Analysis | Values |
| --- | --- |
| **Soil pH** | 5.4 |
| Macronutrients (ppm) |  |
| Phosphorus (P) | 8.0 |
| Potassium (K) | 170.0 |
| Calcium (Ca) | 411.0 |
| Magnesium (Mg) | 78.0 |
| Sulfur (S) | 6.1 |
| Micronutrients (ppm) |  |
| Boron (B) | 0.0 |
| Manganese (Mn) | 3.3 |
| Zinc (Zn) | 4.3 |
| Copper (Cu) | 0.1 |
| Iron (Fe) | 12.3 |
| Aluminum (Al) | 50.0 |
| Lead (Pb) | 6.9 |
| **Cation Exch.** (meg/100 g) | 10.5 |
| **Exch. Acidity** (meg/100 g) | 7.4 |
| **C:N Ratio** | 12.3 |

**Supplementary Table S2.** Taxonomic classification of ASVs using the Greengenes database. ASVs given are the 8 ASVs that contributed the most predictive power to our machine learning model predicting if microbial communities were from hosts under drought or control conditions.

| **ASV ID** | **Greengenes Classification** |
| --- | --- |
| ASV 1 | k__Bacteria;p__Actinobacteria;c__Actinobacteria;o__Actinomycetales;f__;g__;s__ |
| ASV 2 | k__Bacteria;p__Actinobacteria;c__Actinobacteria;o__Actinomycetales;f__Microbacteriaceae;g__Microbacterium;__ |
| ASV 3 | k__Bacteria;p__Actinobacteria;c__Actinobacteria;o__Actinomycetales;f__Microbacteriaceae;g__Frigoribacterium;s__ |
| ASV 4 | k__Bacteria;p__Actinobacteria;c__Actinobacteria;o__Actinomycetales;f__Microbacteriaceae;__;__ |
| ASV 5 | k__Bacteria;p__Actinobacteria;c__Actinobacteria;o__Actinomycetales;f__Beutenbergiaceae;g__Salana;s__multivorans |
| ASV 6 | k__Bacteria;p__Bacteroidetes;c__Sphingobacteriia;o__Sphingobacteriales;f__Sphingobacteriaceae;__;__ |
| ASV 7 | k__Bacteria;p__Proteobacteria;c__Alphaproteobacteria;o__Sphingomonadales;f__Sphingomonadaceae;__;__ |
| ASV 8 | k__Bacteria;p__TM7;c__TM7-3;o__EW055;f__;g__;s__ |

**Supplementary Figures**

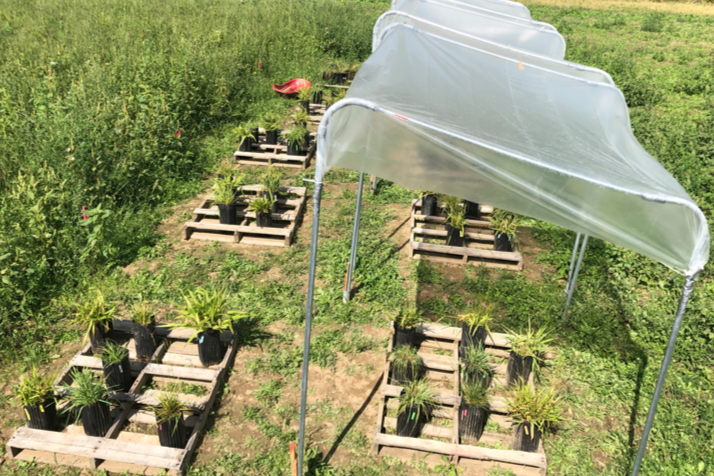

**Supplementary Figure S1.** Pots were organized in a randomized block design. Half the plants were kept under rain shelters to induce drought and the other half were in the open and their soil moisture was maintained above 80% field capacity.

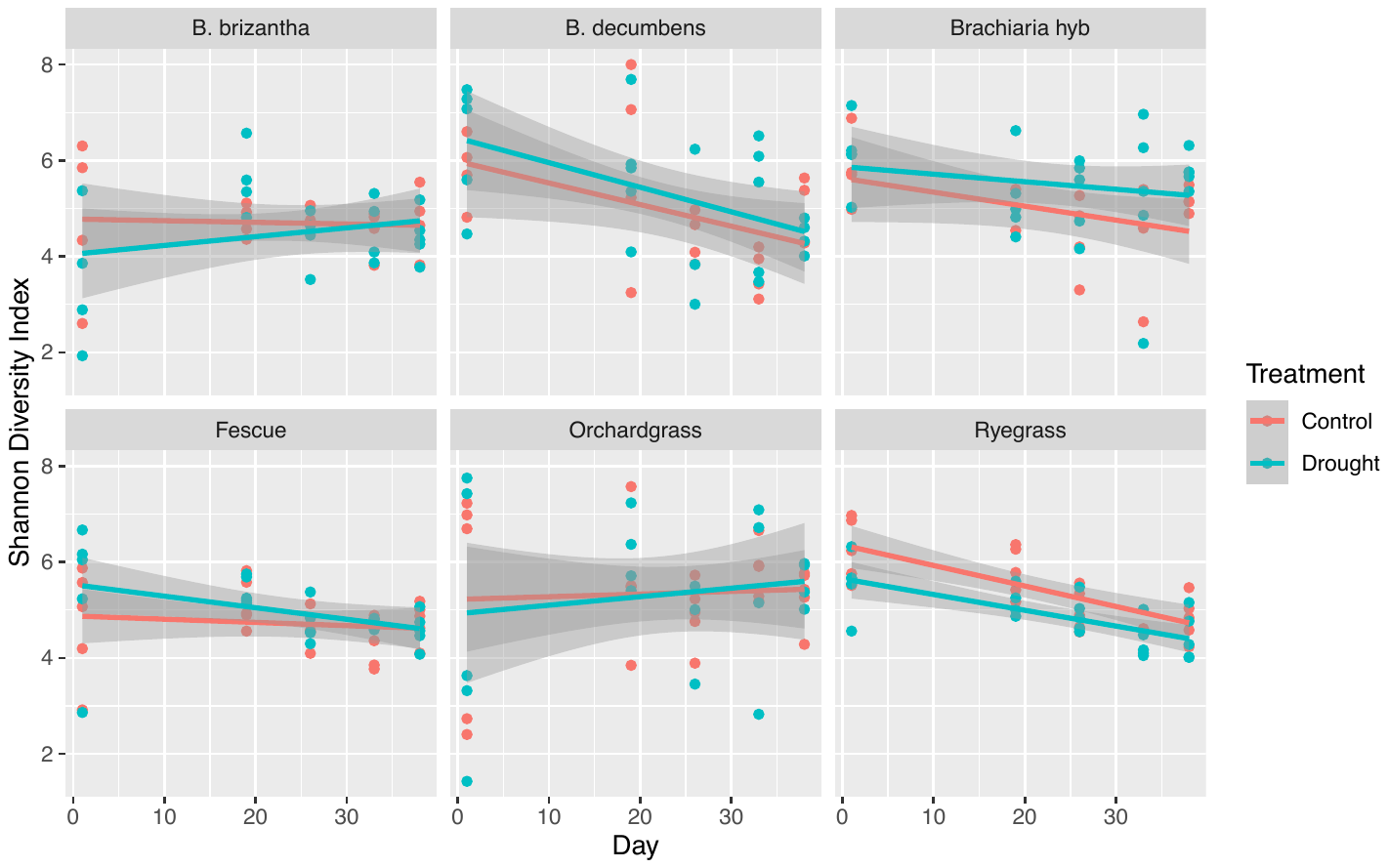

**Supplementary Figure S2.** Alpha diversity of phyllosphere communities did not significantly change as a result of drought stress. Alpha diversity was different between host species and varied over time. Alpha diversity was measured using Shannon Diversity Index, which takes into account both abundance and evenness of species present. We fit an interactive GLMM with treatment, time, and host species as fixed effects and sample ID as a random effect.

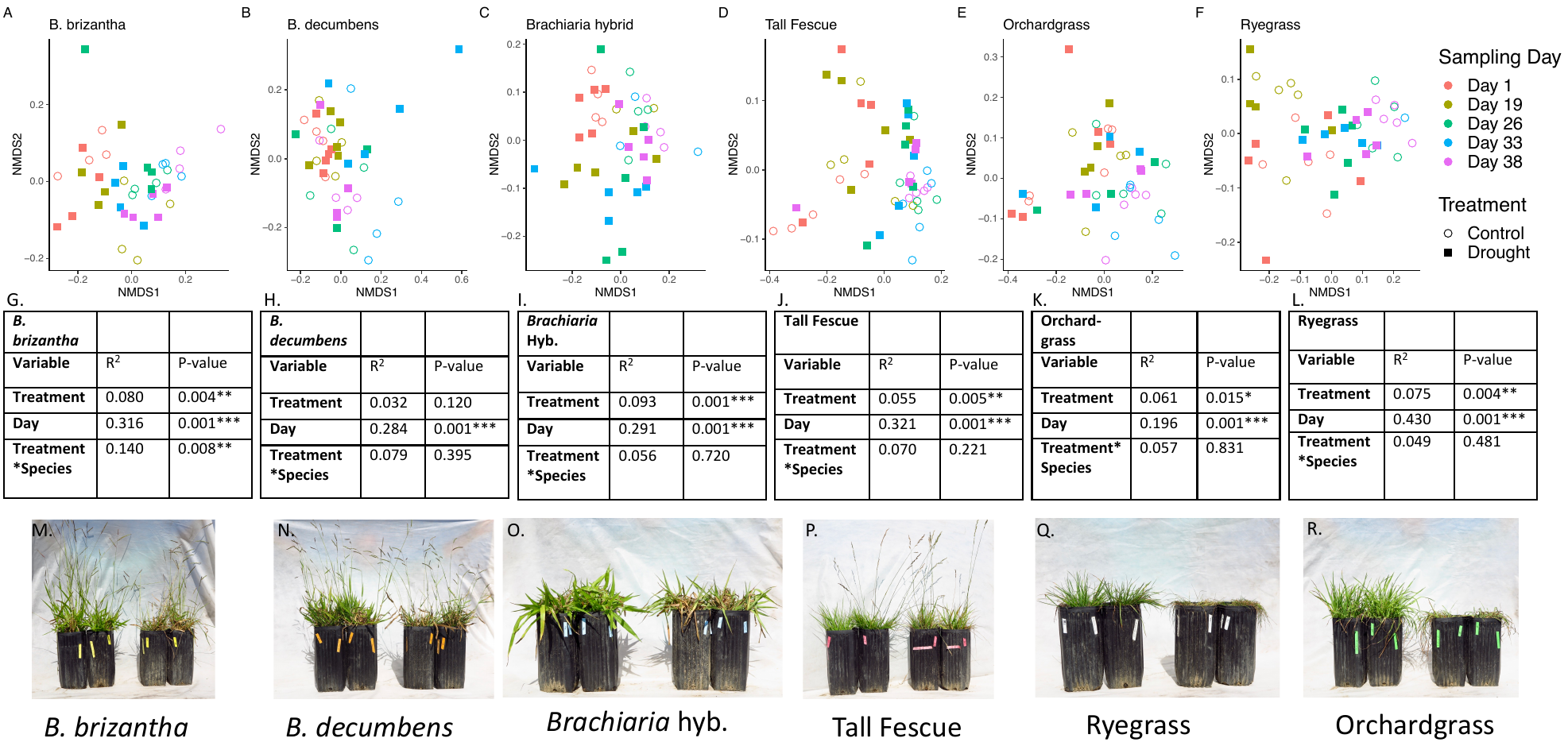

**Supplementary Figure S3.** Bacterial community changes over time and as a result of drought on each species were determined using a PERMANOVA of weighted UniFrac distances (G-L) and visualized using NMDS (A-F). Images of control plants next to plants exposed to drought treatment at the end of the experimental period are given in figures M-R.

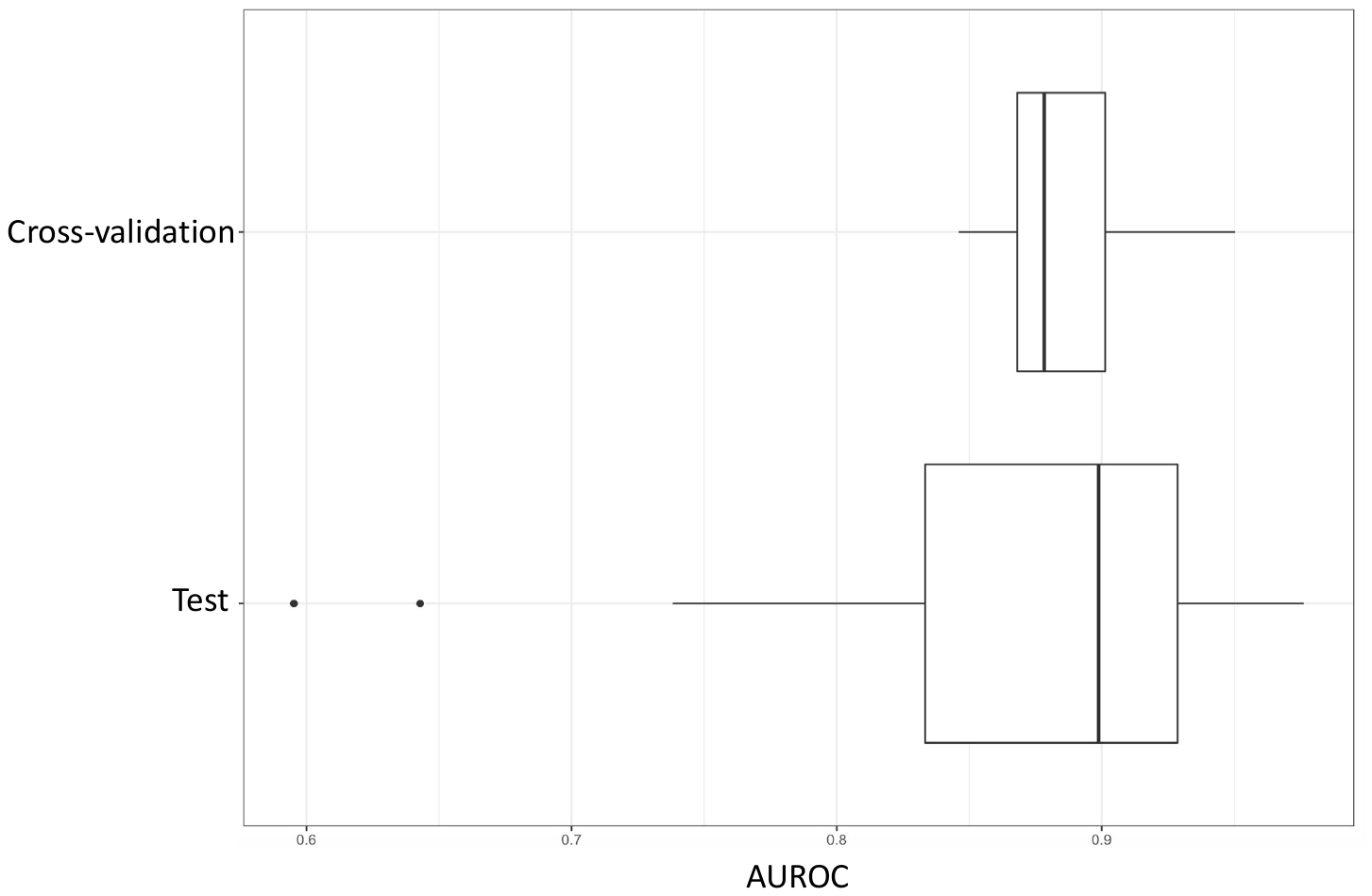

**Supplementary Figure S4.** Machine learning using a random forest model resulted in high predictive power for determining if communities were from control or drought stressed plants. Any points occurring at levels higher than 0.5 in the area under the receiver operating characteristic curve (AUROC) indicates model prediction was better than by chance. Cross-validation and Test data (AUROC of 0.87) resulted in similar AUROC values indicating the model had high predictive power.

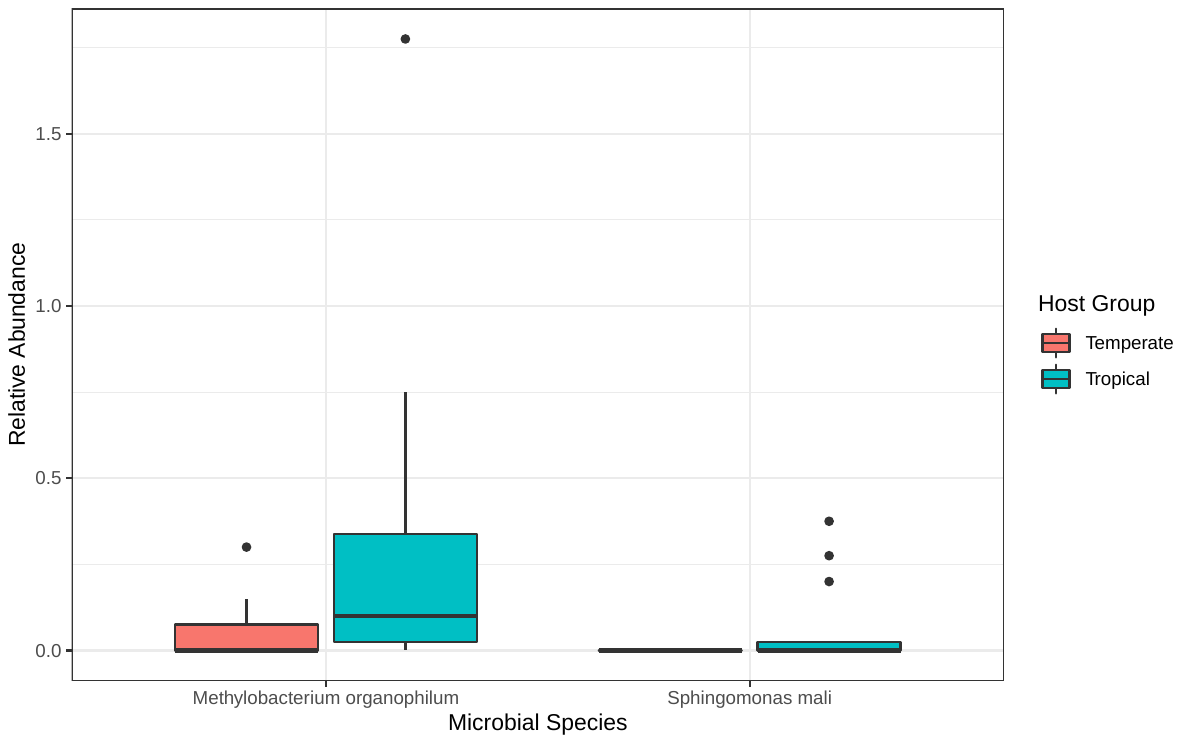

**Supplementary Figure S5**. Two bacterial species were highly important for accurately predicting if a community came from a tropical or temperate host species on the last day of drought. *Methylobacterium organophilum* occurred at a higher relative abundance on tropical grasses compared to temperate grasses. *Sphingomonas mali* was present in low relative abundances on tropical grasses, but it was not detected on any temperate grass host.

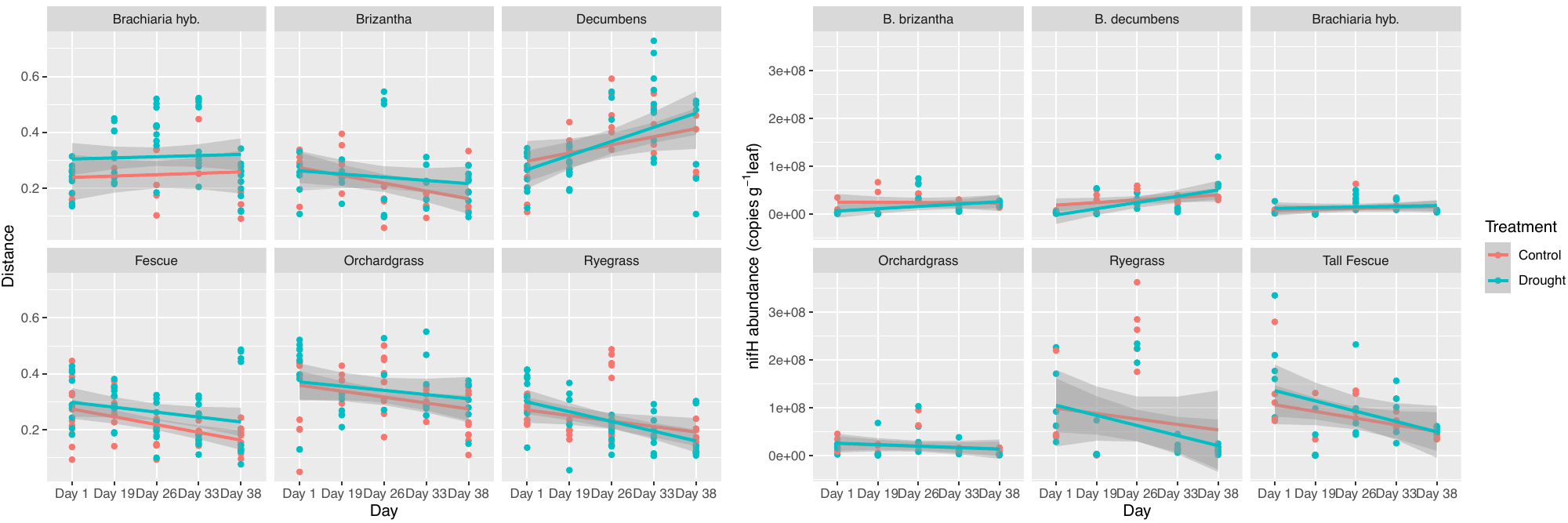

**Supplementary Figure S6.** Changes in average community distance over time measured using weighted UniFrac follows similar trends to changes in *nifH* gene copy abundance.

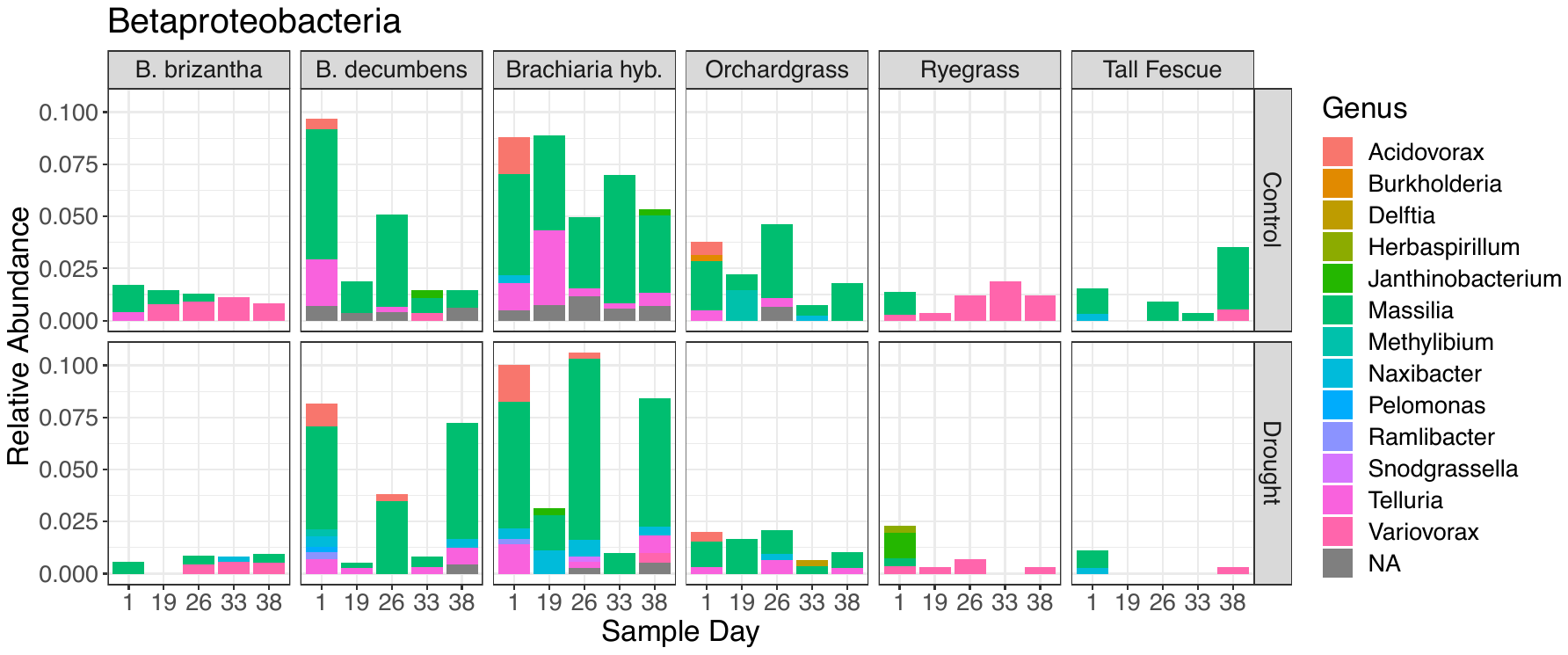

**Supplementary Figure S7.** Average relative abundance of genera belonging to the class Betaproteobacteria over the 38-day experimental period for each control and drought stressed plant host species. The host plant *Brachiaria decumbens* had a significant increase in the genus *Massilia* under severe drought conditions, which is known to have nitrogen fixation potential [4].

**References for Supplementary Information**

1.	Redford AJ, Bowers RM, Knight R, Linhart Y, Fierer N. The ecology of the phyllosphere: geographic and phylogenetic variability in the distribution of bacteria on tree leaves. *Environ Microbiol* 2010; **12**: 2885–2893.

2.	Chelius MK, Triplett EW. The Diversity of Archaea and Bacteria in Association with the Roots of Zea mays L. *Microbial Ecology* 2001; **41**: 252–263.

3.	Poly F, Monrozier LJ, Bally R. Improvement in the RFLP procedure for studying the diversity of nifH genes in communities of nitrogen fixers in soil. *Research in Microbiology* 2001; **152**: 95–103.

4.	Suleiman MK, Quoreshi AM, Bhat NR, Manuvel AJ, Sivadasan MT. Divulging diazotrophic bacterial community structure in Kuwait desert ecosystems and their N2-fixation potential. *PLOS ONE* 2019; **14**: e0220679.
